# Taming the massive genome of Scots pine with PiSy50k, a new genotyping array for conifer research

**DOI:** 10.1101/2021.06.29.450162

**Authors:** Chedly Kastally, Alina K. Niskanen, Annika Perry, Sonja T. Kujala, Komlan Avia, Sandra Cervantes, Matti Haapanen, Robert Kesälahti, Timo A. Kumpula, Tiina M. Mattila, Dario I. Ojeda, Jaakko S. Tyrmi, Witold Wachowiak, Stephen Cavers, Katri Kärkkäinen, Outi Savolainen, Tanja Pyhäjärvi

**Affiliations:** Department of Ecology and Genetics, University of Oulu, 90014 University of Oulu, Finland; UK Centre for Ecology & Hydrology, Bush Estate, Penicuik, Midlothian, UK, EH26 0QB, UK; Natural Resources Institute Finland (Luke), Paavo Havaksen tie 3, 90570 Oulu, Finland; Université de Strasbourg, INRAE, SVQV UMR-A 1131, F-68000, Colmar, France; Natural Resources Institute Finland (Luke), Latokartanonkaari 9, FI-00790 Helsinki, Finland; Department of Organismal Biology, EBC, Uppsala University, Uppsala, Sweden; Norwegian Institute of Bioeconomy Research, Ås, Norway; Institute of Environmental Biology, Faculty of Biology, Adam Mickiewicz University in Poznań, Uniwersytetu Poznańskiego 6, 61-614 Poznań, Poland; Department of Forest Sciences, University of Helsinki, 00014 University of Helsinki, Finland

## Abstract

Scots pine (*Pinus sylvestris*) is the most widespread coniferous tree in the boreal forests of Eurasia and has major economic and ecological importance. However, its large and repetitive genome presents a challenge for conducting genome-wide analyses such as association studies and genomic selection. We present a new 50K SNP genotyping array for Scots pine research, breeding programs, and other applications. To select the SNP set, we first genotyped 480 Scots pine samples on a 407 540 SNP screening array, and identified 47 712 high-quality SNPs for the final array (called ‘PiSy50k’). Here, we provide details of the design and testing, as well as allele frequency estimates from the discovery panel, functional annotation, tissue-specific expression patterns, and expression level information for the SNPs or corresponding genes, when available. We validated the performance of the PiSy50k array using samples from breeding populations from Finland and Scotland. Overall, 39 678 (83.2%) SNPs showed low error rates (mean = 0.92%). Relatedness estimates based on array genotypes were consistent with the expected pedigrees, and the amount of Mendelian error was negligible. In addition, array genotypes successfully discriminate Scots pine populations from different geographic origins. The PiSy50k array will be a valuable tool for future genetic studies and forestry applications.

**Significance statement:** Scots pine is an evolutionary, economically and ecologically impressive coniferous species but its gigantic genome has limited studying e.g. the genetic basis of its functional trait variation. We have developed a genotyping array that facilitates Scots pine genetic research and linking its trait variation to genetic polymorphisms and gene expression levels across the genome.

## Introduction

Scots pine (*Pinus sylvestris*) is one of the world’s most widely distributed conifers (Durrant *et al*., 2016) and is dominant in forests across 145 million hectares in Northern Eurasia (Mason and Aía, 2000; Mullin *et al*., 2011; Pyhäjärvi *et al.*, 2020). The species is an important source of timber and other wood-based products (CABI, 2013) and boreal forests, of which Scots pine is an essential part, are a significant carbon sink (Pan *et al*., 2011). In addition to traditional timber, pulp, paper, and energy production, more diverse uses for Scots pine biomass are currently being developed (e.g., Agbor *et al*., 2011; Rusanen *et al*., 2019). The combination of large biomass volumes, the species capability of adapting to varying marginal environments (Durrant *et al*., 2016), and modern genomic tools provide new possibilities for improving the desired economic and ecological properties.

Breeding activities of Scots pine are centralized in Fennoscandia and the Baltic region, Sweden and Finland having the most advanced breeding programs (Haapanen *et al*., 2015). A first cycle of selection and breeding was completed in the UK in the late 20^th^ century (Lee, 2002), and there is currently substantial interest in further improvement of the species, to reduce national dependency on exotics. The genetic gains from breeding are delivered by seed from seed orchards, comprising copies of field-tested plus trees (outstanding selections from wild stands). Orchard-reproduced stock has been predicted to yield 20-25% improvement in per unit area wood production above unimproved seed lots (Rosvall *et al*., 2001, Haapanen *et al*., 2016; Jansson *et al*., 2017). Forest tree breeding programs traditionally operate on large numbers of individuals. Cost-effective genotyping platforms are therefore essential in incorporating genomics to tree breeding schemes in the extent that is now true for cattle and crop breeding (Grattapaglia *et al*., 2018, Meuwissen *et al*., 2016; Voss-Fels *et al*., 2019).

Genotyping arrays are efficient and easy in comparison to other cost efficient sequencing methods such as genotyping-by-sequencing (Pavan *et al.*, 2020). They are more reproducible across studies, have less missing data and, importantly, require less bioinformatic pre-processing (e.g., Darrier *et al*., 2019). For forest tree species, SNP arrays are available for walnut (Marrano *et al.*, 2019), Norway spruce (Bernhardsson *et al*., 2020) and several eucalypt species (Silva-Junior *et al.*, 2015). They have been used to build linkage maps (Silva-Junior and Grattapaglia, 2015), develop genomic selection (GS) models (Tan *et al.*, 2017) and in genome-wide association studies (GWAS) (Bernard *et al.*, 2020).

We foresee four primary applications for a new Scots pine SNP genotyping array:

### 1) Genomic selection

Genomic selection aims to predict the breeding value of an individual based on its genotypes, where markers are assumed to be in linkage disequilibrium (LD) with the causal variation (Meuwissen et al., 2001). In a set of individuals with both genotype and phenotype data (training population), genomic prediction models are first generated and tested, leading to a prediction equation. Genomic estimated breeding values (GEBV) can then be calculated from this equation for individuals with genotype data only (e.g., Wray *et al*., 2019). GS in trees shows promising results (Isik 2014) and good predictive ability has been achieved with a few thousand of SNPs (e.g., Bartholomé *et al*., 2016; Calleja-Rodriguez *et al*., 2020; Cappa *et al*., 2019; Chen *et al*., 2018; Grattapaglia *et al*., 2018; Lenz *et al*., 2017; Resende *et al*., 2012; but see Thistlethwaite *et al*., 2020).

GS has potential to increase genetic gains per unit of time when the breeding cycle can be shortened, i.e. when reproductive maturity is reached soon after prediction of GEBV. There are significant biological constraints to achieve this in Scots pine that reaches sexual maturity at 8-20 years of age (Sarvas 1964). Nevertheless, genomic markers can provide other benefits by reducing the phenotyping costs and achieving higher selection intensities in situations when a large number of selection candidates are more easily genotyped than phenotyped (Calleja-Rodriguez *et al*., 2020; Grattapaglia *et al*., 2018; Voss-Fels *et al*., 2019). The operational viability of such measures is obviously dependent on the costs of genotyping.

### 2) Pedigree construction

Genotyping data can be used to confirm and reconstruct pedigrees, identify labeling and grafting errors, and estimate genomic relatedness among individuals. Realized genomic relationships are potentially very useful for Scots pine breeding programs, as they allow more accurate genomic prediction of breeding values. Genomic relationships can also help to bridge unconnected progeny-testing series in a multi-environment genetic evaluation. Pedigree reconstruction and parentage analysis using markers also opens opportunities for implementing less costly breeding strategies, such as polymix breeding (Isik 2014).

### 3) Genome-wide association studies

Many of the most valued characteristics of Scots pine and other conifers are complex traits, controlled by many genes. GWAS offers a way to detect the loci responsible governing the variation, improving our understanding of the genetic architecture and biological mechanisms behind these traits (Burghardt *et al.*, 2017; González-Martínez *et al*., 2007 ; Neale and Savolainen, 2004; Yeaman *et al*., 2016). Large sample sizes are crucial for detecting the associations, since polygenic traits are mostly controlled by numerous small effect polymorphisms (Tam *et al.*, 2019; Yang *et al.*, 2010). A genome-wide SNP array is a convenient tool for quickly genotyping many samples. Use of a common genotyping platform will allow for comparison across studies.

### 4) Genetic mapping

High resolution genetic maps inform about the linkage relationships. They are important tools in quantitative trait locus mapping (Lander and Thompson 1990). Combined with physical maps or partial genomic information, they allow analysing of the recombination rate landscape of the genome. To achieve high resolution, large numbers of progeny need to be genotyped, for which SNP arrays are a cost efficient and powerful solution. When SNPs are anchored to scaffolds of a genome assembly, maps derived from SNP array genotyping can be used to improve the scaffolding of reference genomes, by linking together or re-ordering contigs (Fierst, 2015).

In addition, other potential applications for a SNP genotyping array include monitoring genetic diversity, tracing geographic origin, estimating population structure, demographic inference, identifying segregation distortion and identifying large structural variants.

SNP arrays are valuable universal tools for genetic fingerprinting and evaluation of diversity, but they also have limitations. For instance, SNPs are typically accumulated close to or within coding regions, because data are easier to obtain during SNP discovery using RNA-seq or exome-targeted approaches than with whole genome sequences. Further, coding regions are often of high interest and favored in array design. Also, as arrays only score preassigned SNPs with a minimum minor allele frequency (MAF) threshold often applied, there is always an ascertainment bias. This bias affects analyses performed on new individuals using the same set of markers in two ways (McTavish and Hillis, 2015). First, loci with rare alleles in the discovery population will not be scored. This may cause a bias in diversity estimation in favor of those with common alleles. Second, at the population level, allele frequencies, and thus diversity, in samples genetically close to the discovery panel will be biased upward compared to samples from a distant lineage. Ascertainment bias thus is especially problematic for inferences requiring information on rare alleles and not suitable for identifying new genetic variants. However, in many analyses, the ascertainment bias can be taken into account if the original SNP discovery panel and the array design is known (Clark *et al.*, 2005).

Here, we present the Axiom PiSy50k (Thermo Fisher Scientific), a new genotyping array for Scots pine. We describe the different SNP sources and discovery panels and the selection process used during the array design. The final array combines a set of high-performing SNPs from a previously developed Axiom_PineGAP trans-specific SNP array of *Pinus* (Perry *et al.*, 2020) and a new set of curated SNPs originating from exome capture, RNA-seq, PacBio and candidate gene studies (Table 1). We provide a detailed description of SNP discovery, screening, filtering, evaluation of ascertainment bias, error rates, and the metadata we collected during the design, such as gene expression and copy-number variation. We also explore the array’s capability to discriminate populations and reconstruct pedigrees.

**Table 1.**
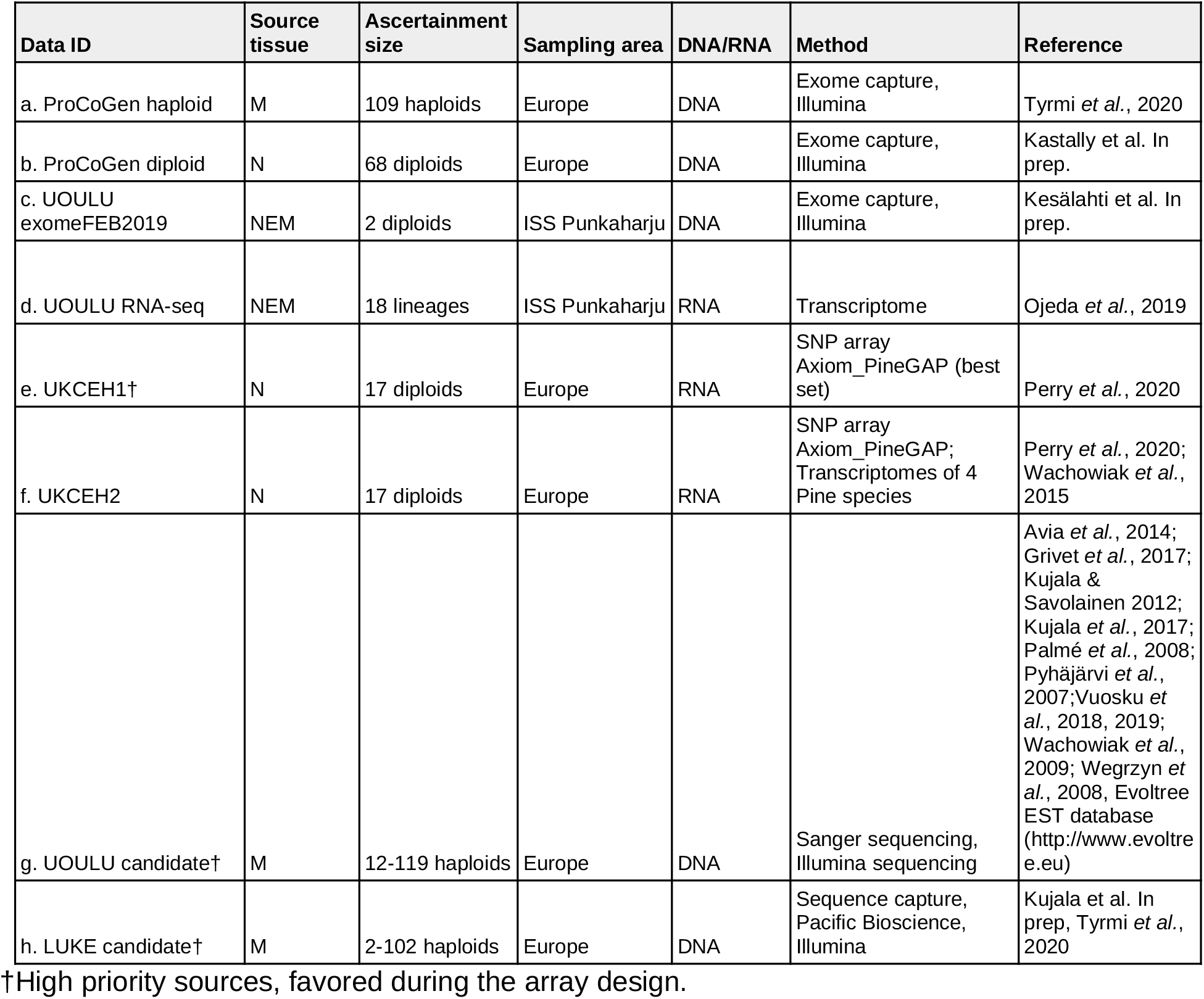
Sources of SNPs used in the design of PiSy50k array (M = megagametophyte, N = needle, E = embryo; ISS Punkaharju = Intensive Study Site Punkaharju in southeast Finland).

| Data ID | Source tissue | Ascertainment size | Sampling area | DNA/RNA | Method | Reference |
| --- | --- | --- | --- | --- | --- | --- |
| a. ProCoGen haploid | M | 109 haploids | Europe | DNA | Exome capture, Illumina | Tyrmi *et al.*, 2020 |
| b. ProCoGen diploid | N | 68 diploids | Europe | DNA | Exome capture, Illumina | Kastally *et al.* In prep. |
| c. UOULU exomeFEB2019 | NEM | 2 diploids | ISS Punkaharju | DNA | Exome capture, Illumina | Kesälähti *et al.* In prep. |
| d. UOULU RNA-seq | NEM | 18 lineages | ISS Punkaharju | RNA | Transcriptome | Ojeda *et al.*, 2019 |
| e. UKCEH1† | N | 17 diploids | Europe | RNA | SNP array<br>Axiom_PineGAP (best set) | Perry *et al.*, 2020 |
| f. UKCEH2 | N | 17 diploids | Europe | RNA | SNP array<br>Axiom_PineGAP;<br>Transcriptomes of 4 Pine species | Perry *et al.*, 2020;<br>Wachowiak *et al.*, 2015 |
| g. UOULU candidate† | M | 12-119 haploids | Europe | DNA | Sanger sequencing,<br>Illumina sequencing | Avia *et al.*, 2014;<br>Grivet *et al.*, 2017;<br>Kujala &<br>Savolainen 2012;<br>Kujala *et al.*, 2017;<br>Palmé *et al.*, 2008;<br>Pyhäjärvi *et al.*, 2007; Vuosku *et al.*, 2018, 2019;<br>Wachowiak *et al.*, 2009; Wegrzyn *et al.*, 2008, Evoltree EST database (http://www.evoltree.eu) |
| h. LUKE candidate† | M | 2-102 haploids | Europe | DNA | Sequence capture,<br>Pacific Bioscience,<br>Illumina | Kujala *et al.* In prep, Tyrmi *et al.*, 2020 |

†High priority sources, favored during the array design.

## Results and discussion

### Array design

SNP choice and array design had four main stages: collection, filtering, *in silico* evaluation and screening array evaluation (Figure 1). We first collected SNPs from eight data sets that differed in sample size, sampling design, source material (RNA or DNA, tissue) and sequencing technology (Sanger sequencing, PacBio, Illumina-seq). We filtered these initial data, tailoring our approach to each data source’s specific characteristics. We removed markers likely to be in paralogous areas of the genome. Paralogy is a common problem for conifer species, which have large genomes with a lot of repetitive elements (Neale *et al.*, 2014). Partly, this was done by checking haplotypes from seed megagametophyte tissue, where observed heterozygosity indicates false SNPs generated by paralogy. After the initial filtering, Thermo Fisher Scientific conducted an *in silico* evaluation of 1.3 million SNPs and from these, we selected 407 540 SNPs of high interest and strong predicted performance.

**Figure 1.**
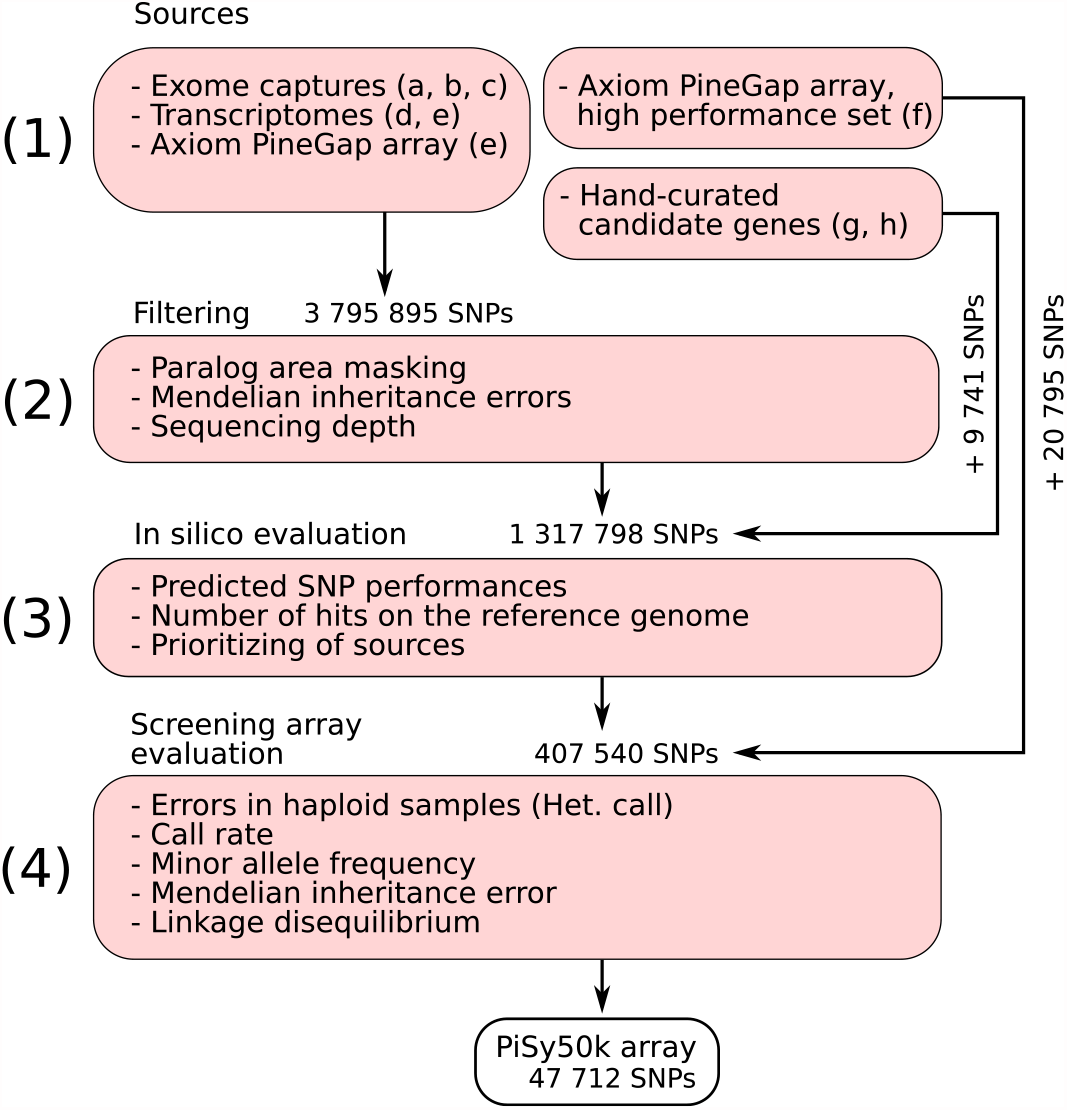
Flow chart of the PiSy50k array design. We proceeded in four steps: (1) the collection of SNPs from 8 sources (Table 1; a: ProCoGen Haploid, b: ProCoGen Diploid, c: UOULU exomeFEB2019, d: UOULU RNA-seq, e: UKCEH2, f: UKCEH1, g: UOULU candidate and h: LUKE PacBio); (2) filtering to remove SNPs from paralogous genomic areas, SNPs with low sequencing depth or Mendelian errors; (3) evaluation to retain the best set of 407 540 markers (screening set) and (4) filtering based on the screening array performance to select the 47 712 markers retained in the PiSy50k array.

### Performance of the screening array

We evaluated the performance of the screening array by genotyping a natural population sample of 470 trees, six megagametophytes and four diploid embryos from full-sib crosses, all from Finland. SNPs were assigned to six classes: Poly High Resolution (PHR, three well-separated genotypesclusters), No Minor Homozygote (NMH, two well-separated genotypes clusters, homozygous and heterozygous), Mono High Resolution (MHR, one homozygous genotype cluster), Call Rate Below Threshold (CRBT), Off-Target Variant (OTV, more than three clusters) and Others. When choosing SNPs for the PiSy50k array based on the screening array, we considered conversion types PHR, NMH and MHR as successful. Of 407 540 SNPs in the screening array, 245 149 (60.2%) converted successfully and 157 325 (38.6%) were polymorphic (Table S1, Figure 2). The success rate varied among sources from 10% to 50%, with lowest and highest rates in the LUKE candidate and UOULU candidate derived SNPs respectively (Table S1, Figure 2). The latter set had already gone through several rounds of verification and thus its higher conversion rate was not surprising. The genotyping success rate at sample level was high; 476 (99%) samples had a call rate above the 97% threshold in the conversion classes PHR, NMH, and MHR.

**Figure 2.**
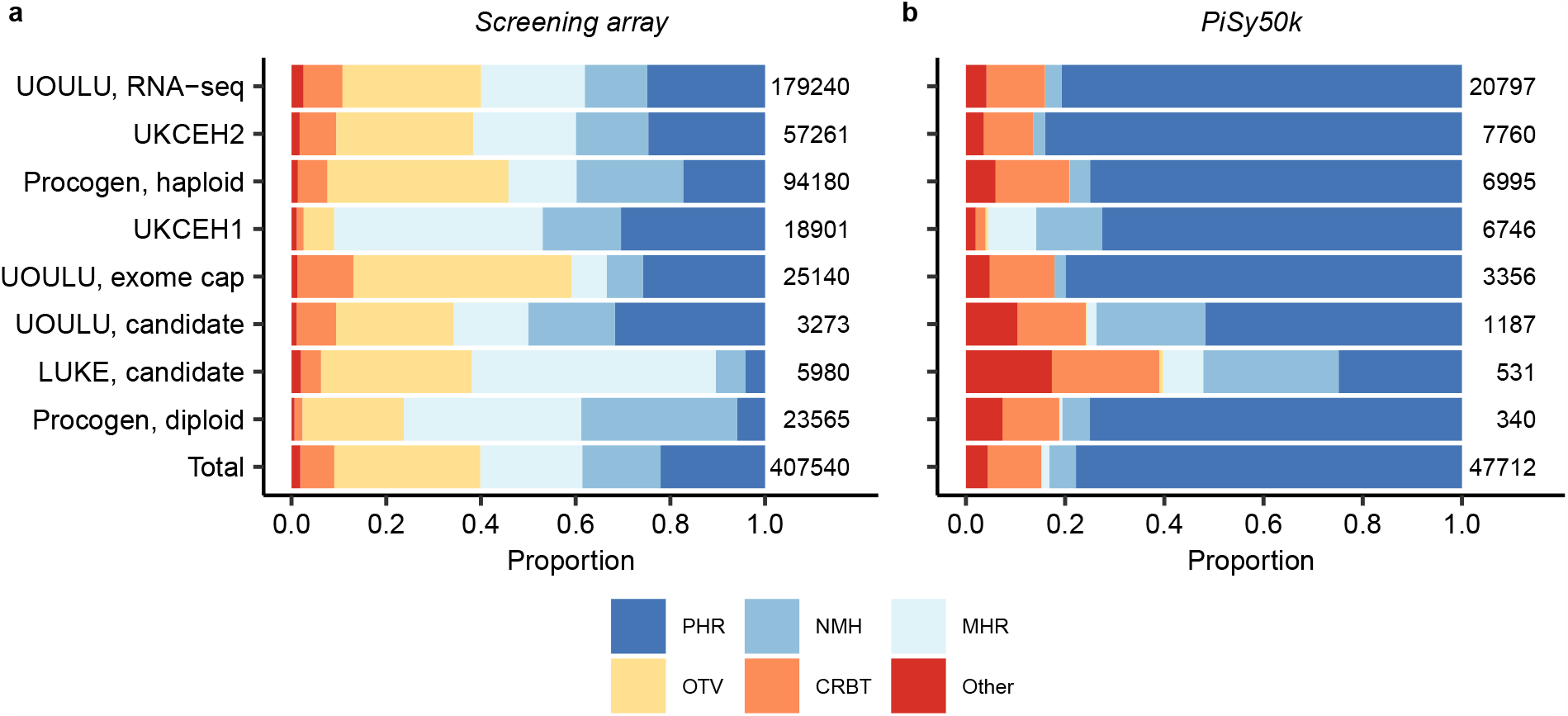
The proportions of conversion types of each marker source in (a) the screening array and (b) the PiSy50k array. PHR = Poly High Resolution, NMH = No Minor Homozygote, MHR = Mono High Resolution, CRBT = Call Rate Below Threshold, OTV = Off-Target Variant. Number right to the bar indicates the total number of SNPs per marker source.

To assess the effects of ascertainment bias throughout the PiSy50k design, we evaluated its effects on the screening array by investigating the minor allele frequency (MAF) distribution and the genetic structure in the sample. The MAF distribution of the screening array is characterized by a deficit of intermediate frequency alleles (MAF values between 0.15 and 0.5) compared to the distribution expected based on the standard neutral model (SNM) (Figure 3A). This is not surprising, as previous studies on Scots pine’s genetic diversity across Europe have demonstrated an overall deficit of intermediate alleles and excess of rare alleles in natural populations of this species compared to the SNM (Tyrmi *et al*., 2020; Pyhäjärvi *et al.*, 2020 and references therein). However, the pattern of rare alleles in the screening set differs from the one in earlier studies. We observed an excess of rare allele classes (MAF between 0.007 and 0.15, Figure S1), but a deficit in the extremely rare classes (MAF below 0.007, Figure S1), as expected from ascertainment bias.

**Figure 3.**
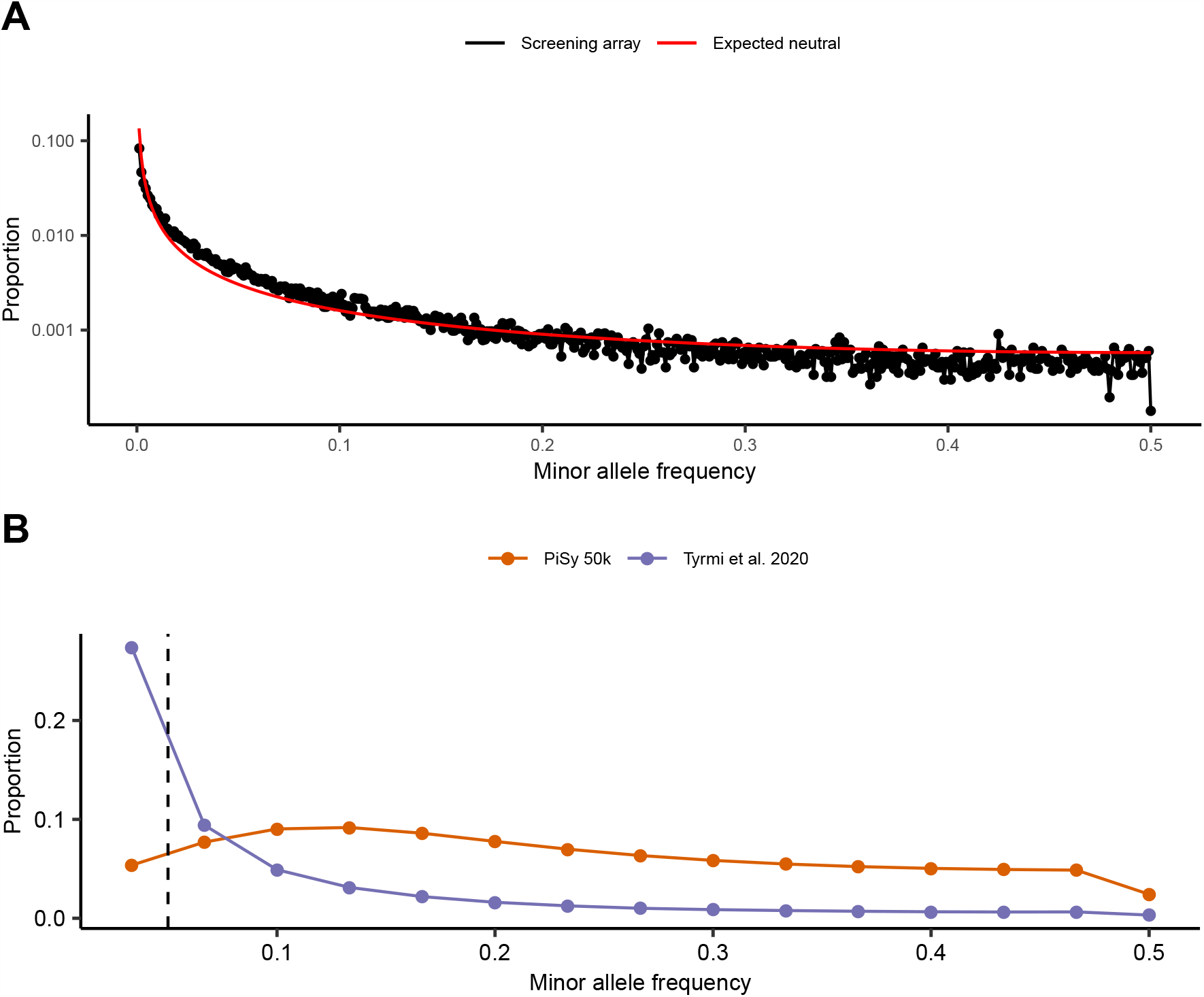
Minor allele frequency (MAF) spectra of the screening and PiSy50k arrays. (A) MAF for the screening population sample (N = 466) and 56 693 SNPs (conversion type PHR and NMR) without missing data in the screening array. The red line illustrates the expected neutral MAF (Tajima, 1989). Note the log scale on the y-axis. (B) MAF based on the PiSy50k array including 38 302 SNPs genotyped in 90 plus-trees across three Finnish breeding populations (red line) and 42 exome captures of Scots pine trees sampled in four natural populations of Finland (Tyrmi et al., 2020). To be comparable, we downsampled both distributions to 30 samples. The vertical dashed line marks the filter threshold of 0.05 used during the array design and below which SNPs were partly excluded. As expected, there is a deficiency of rare alleles in the data obtained from the PiSy50k, as a result of ascertainment bias.

In addition, ascertainment bias influenced the estimates of genetic structure among samples. Principal component (PC) analyses of the screening array genotypes of UOULU RNAseq and UOULU exomeFEB2019 sets clearly separate trees included in the discovery panel from the rest of the samples (Figure S2 a and c). The ascertainment bias was more subtle in the other sources, even when samples from the discovery panel were genotyped (Figure S2 e). This difference was due to the larger size of the other discovery panels (Table 1). The effect of ascertainment bias was particularly severe when the exact discovery panel samples or their close relatives were included (Figure S2 a and c). For most applications and datasets not related to the discovery panels, these effects on genetic structure are unlikely to be so extreme, but we recommend that users of the array carefully consider sample origin when performing analyses.

Finally, from the remaining 75 629 SNPs, we excluded SNPs with heterozygous calls in megagametophyte haploid samples (but allowing one error in SNPs from three high priority sources, see Table 1) or with more than one Mendelian error. We also pruned SNPs in high LD (r² > 0.9), keeping the SNPs with the higher minor allele frequency (MAF) from each such pair. From the remaining loci, we first retained all SNPs from high priority sources and favored SNPs with higher MAF in the remaining set. SNPs in a highly outcrossing wind pollinated natural population of Scots pine are expected to be in Hardy-Weinberg (HW) equilibrium and we used deviation from HW (*p*-values < 0.001) to identify and filter out potentially paralogous and other error prone SNPs. As expected, the markers selected for the PiSy50k array deviated less from the HW expectations and showed less extreme heterozygosities compared to all screening array markers before selection (Figure S3). The final PiSy50k array includes 47 712 SNPs.

### Performance of the PiSy50k array

The 47 712 SNPs in the final PiSy50k array were in 31 657 contigs (average of 1.51 SNPs per contig). Of the eight data sources, markers from RNA-seq origin were the most prevalent (44%; Table S2). The majority of markers have been used in previous studies and come associated with various information depending on the source, including functional annotation, gene expression at the tissue level, and allele frequency estimates in up to 20 European populations (Supporting Data S1).

Altogether, 1 619 markers derived from ProCoGen haploid (1 544) and diploid sources (75) were located on one of the 4 226 scaffolds mapped on *P. taeda* linkage map (Westbrook *et al*., 2015; Figure 4, Table S3). There was an average of 134 SNPs per linkage group (LG) and they were homogeneously distributed among LGs. Even though the majority of SNPs do not have a known position on the map yet, the quick genotyping of large numbers of progeny with the PiSy50k array could be used to improve the genetic map of Scots pine and help anchor genomic reads, scaffolds and SNPs at the chromosome scale in the future.

**Figure 4.**
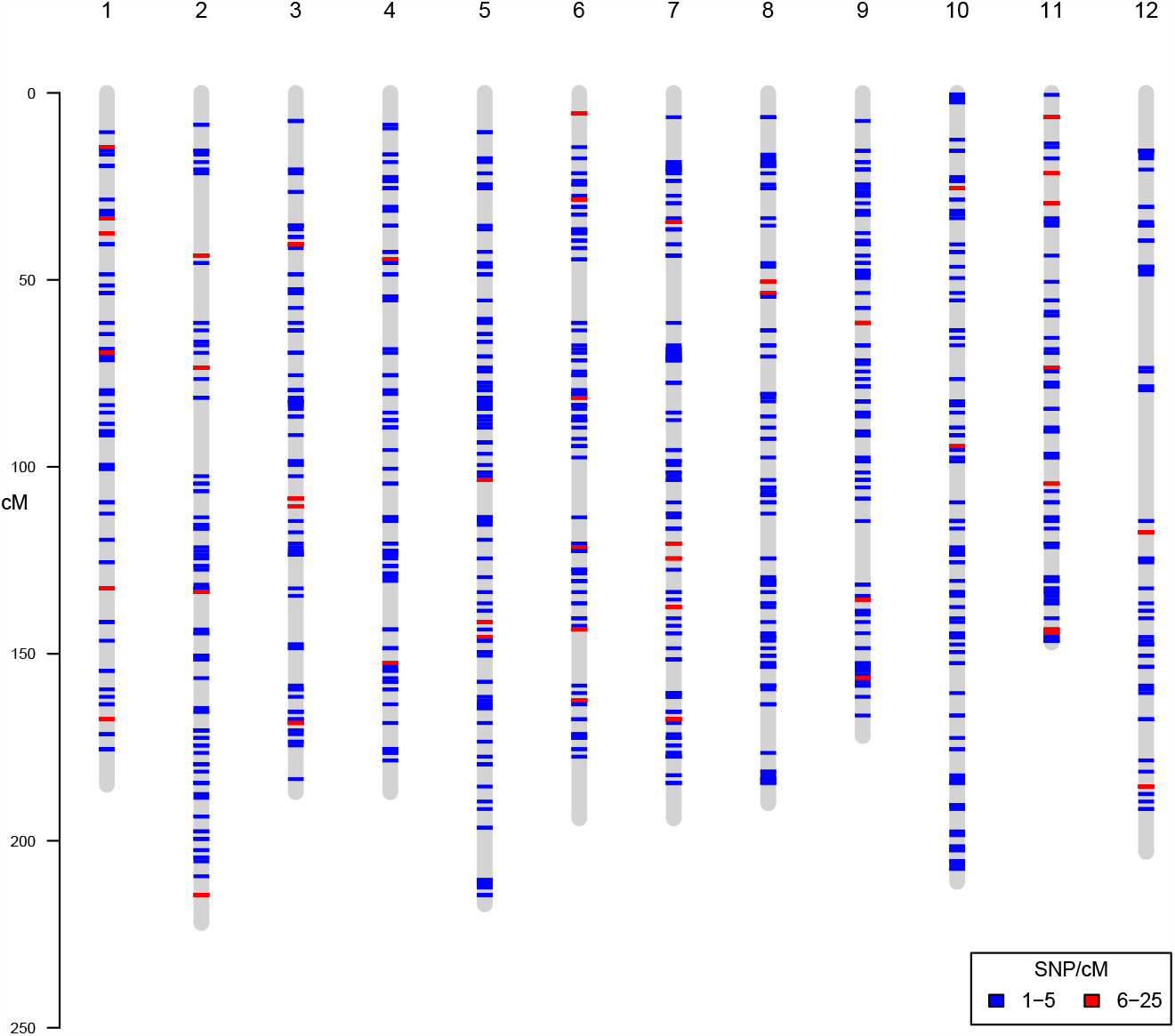
Position and density of 1 619 SNPs from the PiSy50k array on the *P. taeda* linkage map (Westbrook et al., 2015). The vertical grey lines represent the 12 linkage groups in *P. taeda*, while horizontal colored lines indicate the marker positions and density. This plot was made with the R package chromPlot (v 1.12.0) (Oróstica and Verdugo, 2016).

We evaluated the performance of the PiSy50k array by genotyping 2 688 samples from Finland (2178, including 14 controls), Scotland (496), Australia (3), and Estonia (11). Of these, 2 308 samples had call rates above 97% (85.9% of samples), the recommended threshold for Axiom genotyping arrays. In total, 40 405 (84.69 %) markers were successfully converted of which 39 678 markers were polymorphic (Table S4).

Of the 21 control samples, three needle and six megagametophyte samples passed the 97% CR threshold (Table S5). Of the six megagametophyte samples, one replicated pair was recovered. Based on the five control samples retained (three needles and one megagametophyte pair), the error rates were relatively low (mean 0.83%). The error rate in the subset of SNPs shared with the Axiom_PineGAP suggests a similar, or slightly lower, error rate in the PiSy50k (mean 0.52% compared to 0.64% in the Axiom_PineGAP). Overall, these values are close to those obtained in other arrays, e.g. 0.8% in the walnut genotyping array (Marrano *et al.*, 2019), 0.1% in Affymetrix GeneChip Human mapping 50k Array (Saunders *et al.*, 2007), or ranging between 0.03% and 0.05% in the Axiom Apple480K genotyping array (Bianco *et al.*, 2016).

Of the 930 markers with errors among pairs (including both needle and megagametophyte controls), the majority (N = 916) were not shared among controls. This suggests that the error probably occurred during the genotype call for a single sample only, as opposed to the marker itself being unreliable. There are 14 markers for which errors were observed among both megagametophyte and needle controls and they are indicated in Supporting Data S2. Comparison of markers shared between the PiSy50k and Axiom_PineGAP arrays (N = 7592) using the needle control present on both arrays also showed low error rates (mean 0.55%, Table S5) indicating cross-array reproducibility, which allows data obtained by the two arrays to be combined.

To confirm that the variants at the selected SNPs in the PiSy50k array are indeed allelic (not paralog), we assessed the heterozygosity levels of the megagametophyte samples. The two megagametophyte replicates have very low heterozygosity levels (mean 0.89%) compared to the needle replicates (mean 29.30%), suggesting a low level of errors due to paralogy. Of the 40 405 converted markers, 38 906 were homozygous in both replicates, 1 060 were ‘no call’ in at least one replicate, 165 were heterozygous in both replicates and 274 were homozygous in one replicate and heterozygous in the other. The SNPs that were heterozygous in the megagametophyte samples are indicated in Supporting Data S2.

To evaluate the potential of the PiSy50k array for pedigree reconstruction and assess the proportion of Mendelian errors in the array, we analyzed the pairwise relatedness of the full-sib progeny and their parents in a subset of 135 trios across 10 families of our sample. By plotting the kinship coefficient (K, (Manichaikul *et al*., 2010)) against the proportion of sites where individuals share no allele (IBS0), we identified four distinct groups (Figure 5a): (1) known parent-offspring pairs (mean +/- standard deviations: K = 0.245 +/- 0.004, IBS0 = 0.001 +/- 2e-04), (2) full-sibs (K = 0.246 +/- 0.027, IBS0 = 0.015 +/- 4e-03), (3) half-sibs (K = 0.120 +/- 0.018, IBS0 = 0.030 +/- 4e-03), and finally (4) the remaining unrelated pairs (K = -0.002 +/- 0.009; IBS0 = 0.059 +/- 2e-03). We separated parent-offspring pairs from full-sibs, which have expected K values close to 0.25, using the IBS0 statistic (equal or close to 0 between a parent and an offspring but with higher values between siblings (Manichaikul *et al.*, 2010)). Within each family, the K estimates were around the expected value of 0.25, while between families it was close to 0, except for progeny pairs between families 5 and 31, and families 14 and 20, which shared a common parent and had a K estimate around 0.125, as expected for half-sibs (Figure 5). The pedigree relationships identified with PiSy50k matched those expected from the crossing design, demonstrating the array’s power to resolve relatedness structure and reconstruct pedigrees, a critical feature for a multitude of applications in tree breeding and genetics: GWAS, GS, breeding program management and seed production.

**Figure 5.**
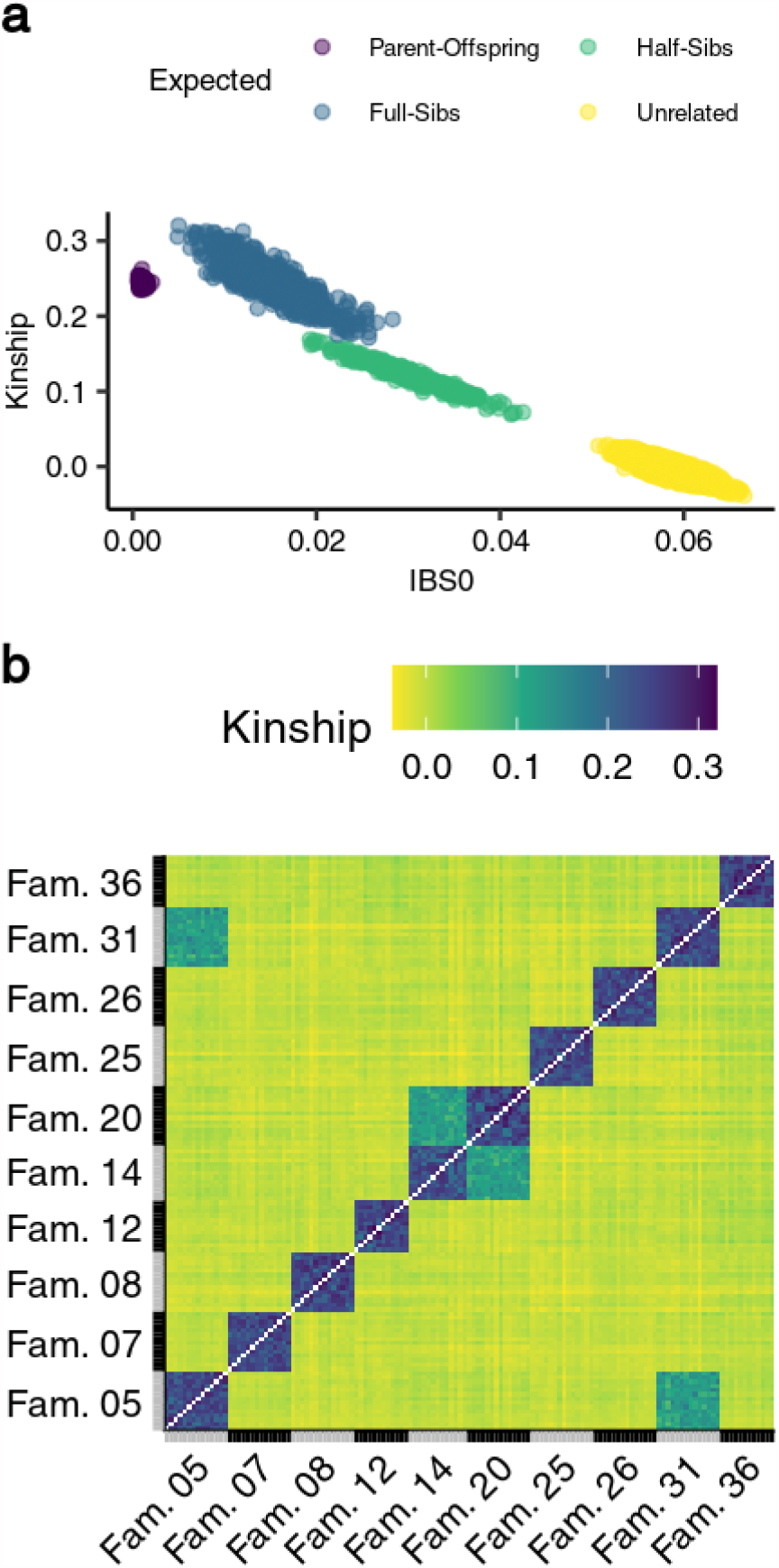
Relatedness analyses of 10 families (including 18 parents and 135 offspring) using the PiSy50k array. (a) Kinship coefficients (Manichaikul et al., 2010) and proportion of sites where individuals share no allele (IBS0) between all pairs and using 39 678 SNPs (PHR + NMH). Expected relationships between pairs are outlined: parent-offspring in purple, full sibs in blue, half sibs in green, and unrelated pairs in yellow. (b) Heat map of the kinship coefficients between all pairs of the 135 offspring.

To further assess the error rate in the PiSy50k data, we evaluated the number of Mendelian errors (ME) within each family. We examined all 40 405 SNPs in 135 trios and identified 16 040 errors across 5 837 loci (mean error rate per locus = 0.29%; Figure S4a). More than 98% of all SNPs had a ME below 5%. Across families, we identified an average of 1 604 errors per family, majority in different SNPs across families (Figure S4b: 4277 SNPs with an error only in a single family and 1110 in at least two). These values are in line with the ME measured in other arrays (Bernhardsson *et al*., 2020; Silva-Junior *et al*., 2015).

### Genetic diversity

To explore the power of genotypes from the PiSy50k array to discriminate trees from different geographic origins, we ran a principal component analysis (PCA) using a subset of 120 samples from different localities in Scotland and Finland (Figure 6). The first two PCs separated two main groups consistent with the two countries of origin. We then ran PCAs using only samples from each country. Although no distinct groups appeared in those analyses, some differentiation was found between samples from different geographic origins in Scotland (Figure 6b) – a level of geographic resolution not previously possible. In the Finnish subset, variation was more homogeneous with less geographic structure (Figure 6c), although samples from Northern origins were located slightly apart from samples from Southern and Central origins.

**Figure 6.**
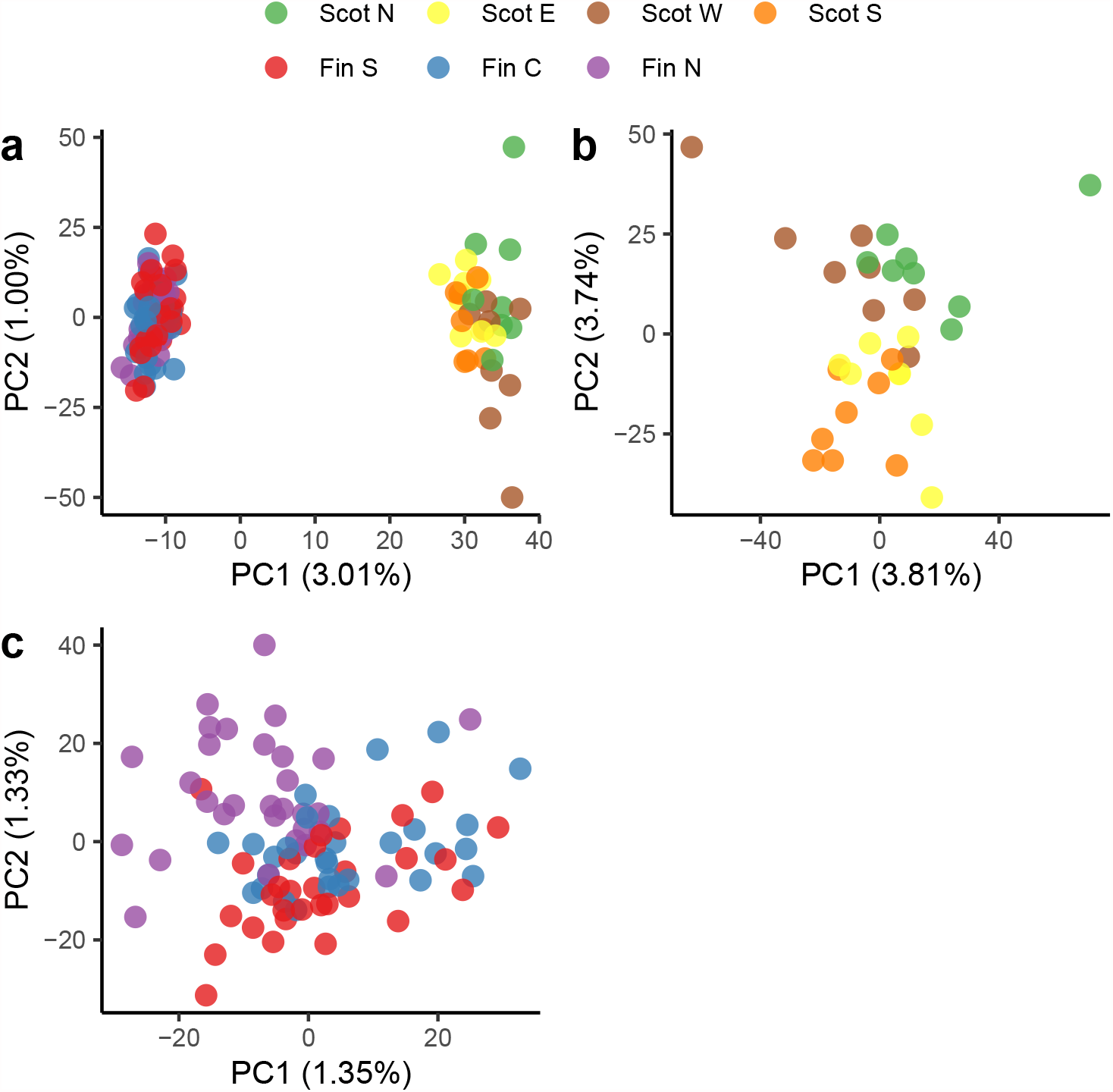
Principal Component Analysis (PCA) using 39 678 polymorphic SNPs from the PiSy50k array genotyped in 120 trees from seven areas in Finland (90) and Scotland y (30). PCA including (a) all 120 samples from Finland and Scotland, (b) 30 samples collected across 21 localities grouped in four geographical areas of Scotland, or (c) 90 samples from Southern, Central and Northern Finland (30 samples each). Scot N, E, W and S: Northern, Eastern, Western and Southern Scotland. Fin S, C and N: Southern, Central and Northern Finland.

To assess the effects of ascertainment bias on the MAF distribution in the PiSy50k array, we compared the frequency distributions obtained from the array to a previously published exome capture dataset (Tyrmi *et al*., 2020) (Figure 3B). We observed a similar but stronger effect of ascertainment on the MAF estimated with the PiSy50k array genotyping results than with the screening array results. Indeed, in the PiSy50k results, the distribution reaches a maximum at frequency 0.13, with decreasing frequencies of lower MAF values, as opposed to the screening array where the peak is at the lowest allele frequency class. This could be explained by the more stringent filtering of SNPs with low allele frequencies when selecting markers for the final PiSy50k set, whereas there was no intentional allele frequency filtering from the source data to the screening set. In addition, the discovery process naturally has an inherent filter for allele frequency, which is the sample size of the discovery panel.

In summary, PiSy50k is a novel genotyping tool for Scots pine, an economically important and widely distributed conifer. It greatly improves the genotyping capacity for the species, which will facilitate wide application of modern breeding tools and supports the development of a new, forest-based bioeconomy. The metadata provided connects the genotyping data to functional properties via annotations and tissue-specific expression patterns. Low error rates indicate high reproducibility even across the previous Scots pine array Axiom_PineGAP (Perry *et al*., 2020), hence new datasets will be back-compatible and all new work will add value to our knowledge of the species.

## Experimental procedure

### Selection of SNPs for initial screening

#### ProCoGen haploid and diploid sets

The ProCoGen haploid and diploid sets were generated with two exome-capture experiments both based on the same bait set used by Tyrmi *et al*. (2020). A total of 177 trees collected across Europe, from Spain to northern Finland, were genotyped using DNA extracted from megagametophyte tissue (haploid set, 109 samples, 12 populations) or needles (diploid set, 68 samples, 8 populations). Bait design, DNA extraction, library preparation, and sequencing steps followed the procedure described in (Tyrmi *et al.*, 2020). We processed the sequences generated to identify SNPs following the same method described in Tyrmi *et al*. (2020) for the haploid set, but applied a few adjustments for the diploid set: we used BWA (Li, 2013) for mapping reads and used samtools v0.9 (command *mpileup*, default parameters) (Li *et al*., 2009) for variant calling. To filter potential paralogs, we removed loci with heterozygous calls in the haploid set or significantly departing from the HWE in the diploid set (PLINK v1.90b5.2 (Chang *et al.*, 2015) command --hardy, at alpha = 0.05). During this procedure, we excluded one haploid sample with an exceptionally high proportion of heterozygous calls. Finally, we excluded all SNPs within 50 bp distance of these markers. We retained 248 591 and 32 649 SNPs in the haploid and diploid sets, respectively.

#### UOULU exomeFEB2019

We used 95 504 SNPs identified in exome capture of a family originating from Punkaharju ISS, in southeast Finland: a cross between Maternal tree 463 and paternal tree 485 (Kesälahti *et al*., In Prep). The material sampled consisted of: needles of both parental trees, one megagametophyte of the paternal tree, two megagametophyte of the maternal tree from open-pollinated seeds, and, from two seeds of the cross progeny, two embryos and a megagametophyte were sampled. We excluded positions with depth below 4 per genotype. We removed twenty-five base pairs both upstream and downstream from each heterozygous site found in haploid megagametophyte as potential areas with paralog or mapping issues.

#### UOULU RNA-seq

The UOULU RNA-seq set refers to markers derived from RNA-seq data (Ojeda *et al.*, 2019) originating from five tissues (needle, phloem, vegetative bud, embryo and megagametophyte) of six unrelated individuals of Scots pine (but 18 haploid genomes when accounting for diploidy and paternal contribution in embryos) collected from Punkaharju ISS. We considered 1 349 291 SNPs obtained by mapping RNA-seq reads to the Scots pine reference transcriptome (https://a3s.fi/pinus_sylvestris_transcriptome_public_/Trinity_CD-HIT.fa). From this initial set, we first excluded markers identified in contigs associated with potential contaminants (fungi or microbes) (Cervantes *et al.* ; Ojeda *et al.*, 2019) (https://a3s.fi/pinus_sylvestris_transcriptome_public_data/Trinity_guided_gene_level_info.txt). Second, we removed heterozygous SNPs in haploid samples. Finally, we compared the genotypes called in megagametophyte, embryo and diploid tissues collected from the same tree to identify and exclude loci with Mendelian errors. In total, we retained 736 827 SNPs.

For the *UOULU RNA-seq* set, we provide information about the predicted multi-copy status, orthologous genes identified in *P. taeda* (Zimin *et al.*, 2014) *and P. lambertiana* (Stevens *et al.*, 2016) based on blastn results (see details in Ojeda *et al.*, 2019), and expression levels and tissue-specificity in five tissues (Cervantes *et al.*). This information is available in Supporting Data S1.

#### UOULU candidate

The UOULU candidate set contains SNPs reported in multiple publications and genetic databases on various candidate genes of Scots pine. This set includes SNP markers used in Kujala *et al*. (2017), and additional SNPs from phenology related genes (Kujala and Savolainen 2012; Palme *et al*., 2008; Pyhäjärvi *et al*., 2007, Wachowiak *et al*., 2009), stress and phenology related genes (Avia *et al.*, 2014), polyamine genes (Vuosku *et al.*, 2018, 2019), genes from comparative resequencing projects (Wegrzyn *et al.*, 2008 ; Grivet *et al*., 2017), and markers identified in sequences from the Evoltree EST database (www.evoltree.eu). Additionally, for a subset of those markers, we have collected allele frequency estimates from two genotyping assay experiments on 426 Scots pine trees (data unpublished). These SNPs, referred to as UOULU candidate VIP in the metadata, were given higher priority during the array manufacture, in both the screening and PiSy50k arrays, by increasing their probeset counts and, this way, improving their call rates during the genotyping.

#### LUKE candidate

The LUKE candidate set comprises SNPs extracted from candidate genes related to phenology (e.g. Bouché *et al*., 2016) and genes of the primary and secondary metabolism pathways active during heartwood formation (Lim *et al.*, 2016). DNA libraries targeting these candidate genes were produced from one individual of Southern Finnish origin and sequenced using a PacBio sequencer (Kujala *et al*., in prep). We used the long PacBio sequences as a reference to map short reads from exome captures of megagametophyte samples of Scots pine collected across Europe (Tyrmi *et al*., (2020) excluding samples from Baza, Spain) with BWA mem (Li, 2013). Since a preliminary variant calling based on this initial mapping resulted in a large number of errors (heterozygous calls in haploid samples), we isolated short reads mapping to individual PacBio contigs and re-assembled them with MIRA (Chevreux, 2007) for each individual. We then aligned the resulting individual re-assemblies to each other with cap3 (Huang and Madan, 1999), and called variants using bcftools (commands mpileup and call). In addition, some SNPs were identified and included solely as being polymorphic within the reference individual.

#### UKCEH sets 1 and 2

We used SNPs collected during the Axiom_PineGAP (Thermo Fisher Scientific) array design (Perry *et al.*, 2020) and from the comparative transcriptomics of four pine species (*P. sylvestris, P. mugo, P. uncinata* and *P. uliginosa*) by Wachowiak *et al*. (2015). Briefly, we identified 196 636 polymorphic positions from transcriptomes, candidate gene sequences and markers from previous population genetic studies on the four above mentioned pine species. From these, we retained two distinct sets: (1) UKCEH1, comprised of 20 795 successfully converted SNPs from the Axiom_PineGAP array, and (2) UKCEH2, a set of 175 841 SNPs including 29 034 SNPs from the Axiom_PineGAP array which were not successfully converted, 31 897 SNPs that passed the initial filtering during the design but were not included in the final array and 114 910 SNPs identified by Wachowiak et al. (2015) which were polymorphic in Scots pine but not included in the Axiom_PineGAP array design.

### SNP scoring for inclusion in the screening array

For each retained site, we built 71-mer probes by extracting up to 35 bp up- and downstream from the source references. We submitted 1 317 798 probes to Microarray Research Services Laboratory (Thermo Fisher Scientific), Santa Clara, US, for scoring (Table S1). During this step, probes’ score were downgraded if: they contained polymorphic sites within 35 bp distance of the focal marker (interfering polymorphism), they were mapped to highly repetitive regions of the genome (using TrinityCD-HIT.fasta.gz and Pita 1.01 (https://treegenesdb.org/FTP/Genomes/Pita/v1.01/genome/Pita.1_01.fa.gz) as references for RNA and DNA based probes respectively), or were highly similar to other probes. Each marker was given a classification: ‘Recommended’, ‘Neutral’, ‘Not recommended’ or ‘Not possible’.

Based on Thermo Fisher Scientific’s evaluation and the available metadata on each data source, we established the following priority groups (in order of priority): (1) the 20 795 high quality SNPs from the Axiom_PineGAP array, (2) all recommended or neutral markers identified by Thermo Fisher Scientific, (3) UOULU candidate markers, (4) LUKE candidate markers, (5) from the ‘not recommended’ set in the ProCoGen haploid set, SNPs of high interest identified in (Tyrmi *et al*., 2020), (6) SNPs with less than 50% of missing data in the discovery panel from the ‘not recommended’ set in the ProCoGen sets, and finally, (7) we relaxed the filtering criterion used by Thermo Fisher Scientific and selected the best markers in the remaining set. More specifically, we relaxed the wobble count filter threshold (number of polymorphic sites on the same 71-mer) from < 4 to < 6, based on the assumption that a high proportion of the variable sites are associated with rare alleles, and thus interfering polymorphism should have lower impact on the probe performance in the case of Scots pine. During the screening array manufacture, out of the 428 516 SNPs retained, a total of 407 540 markers were fitted on the array.

### Screening set genotyping

The screening set of 407 540 SNPs was used to confirm the normal segregation of polymorphism in a larger sample from a natural population, to identify potential deviations from HW equilibrium, indications of paralog mapping — such as heterozygote sites in haploid samples, deviations from Mendelian segregation, and identification of loci in strong LD with each other. To this end, we used the screening array to genotype 480 samples of Scots pine from Punkaharju ISS population, including: 470 diploid needle samples from adult trees, six haploid megagametophytes and four diploid embryos. Two families, “ 463 × 485” and “ 320 ⨯ 251”, with two parents and two offspring (embryos) from each were used to estimate Mendelian error rate.

DNA was extracted from dry needles and fresh megagametophytes using E.Z.N.A.® SP Plant DNA Kit (Omega Bio-tek, Inc.). Genotyping and array manufacturing for the screening set was performed by Thermo Fisher Scientific at Santa Clara, US. Genotype calling was performed by Thermo Fisher Scientific (Applied Biosystems^™^ Axiom^™^ Genotyping Services) following the Axiom Best Practices Workflow (Axiom Genotyping Solution Data Analysis Guide). In short, genotype clusters were defined using samples with quality control call rate (QC CR) >= 0.97 and dish quality control rate (dQC) >= 0.82. The markers were classified into five conversion categories: PolyHighResolution (PHR), NoMinorHom (NMH), MonoHighResolution (MHR), CallRateBelowThreshold (CRBT), Off-Target Variant (OTV), and Other. We retained markers only from classes PHR and NMH with call rate (CR) >= 0.97 in the subsequent analyses of the screening array and for inclusion on the PiSy50k array.

During the array design of both screening and PiSy50k arrays, identical SNPs discovered independently across different sources were identified and merged. To keep track of as much information as possible for those markers, we recorded their common presence and IDs in different sources but eventually assigned a single authoritative origin.

### Selection of markers for the PiSy50k array

For the PiSy50k array, we filtered the markers based on their performance on the screening array prioritizing markers in candidate genes of interest or markers that performed well in the Axiom_PineGAP array (Perry et al., 2020). These markers were within the Axiom Best Practices Workflow default quality thresholds (see above). For each marker with conversion type PHR or NMH, we estimated MAF and tested departure from HW equilibrium (exact test) for 466 individuals, excluding the haploid megagametophyte samples, the offspring samples and four samples with QC CR < 0.97 using PLINK version 1.9 (Purcell et al., 2007). We estimated the number of Mendelian errors in PLINK using the family data.

We excluded markers deviating from HW equilibrium (p < 0.001) and markers with more than one Mendelian error. Markers from the candidate gene sources (LUKE candidate and UOULU candidate) were selected using a lenient inclusion threshold of MAF >= 0.01 and marker call rate > 0.90, which also included markers from the Thermo Fisher Scientific conversion type “ call rate below threshold”. We filtered SNPs from the Axiom_PineGAP array first to include markers with MAF >= 0.05. To increase the number of well performing markers, we also included markers with MAF >= 0.05 in previously genotyped European samples (Perry *et al*. 2020).

To avoid markers in paralogous genomic regions, we excluded markers with heterozygous call in the haploid megagametophyte samples except in three high priority sources (UKCEH1, LUKE candidate and UOULU candidate) for which we allowed at most one, erroneous, heterozygous call per marker. We further granted 381 markers of high interest from sources UOULU candidate (335) and UOULU RNA-seq (23) a higher probeset count in the array to increase their call rate. Finally, to remove the excess from the retained set, we excluded markers from the low priority sources with lowest MAF (MAF after filtering >= 0.08). The final number of markers for PiSy50k was 47 712 (Figure 1). The distribution of the markers by source is shown in Table S2. To inspect how SNP selection for the PiSy50k array affected HW deviation compared to the screening array on average, we plotted the observed *p*-alues from the exact HW tests against the expected *p*-values based on the null distribution in a cumulative Q-Q plot before and after SNP selection. We compared the observed *p*-values of 10 000 random loci against 100 samples drawn from the null distribution using HardyWeinberg package (Graffelman, 2015) in R (The R Project for Statistical Computing)(version 3.6.3). We also illustrated the distribution of genotypes with respect to HW expectations in ternary plots showing genotypes before and after the PiSy50k SNP choice.

To assess the effects of ascertainment bias on the screening array, we ran two analyses. First, we plotted the MAF distribution for loci with conversion types PHR or NMH (n loci without missing data = 56 693, n individuals = 466) against the expected MAF assuming a standard neutral model (Tajima, 1989). Second, we looked at the effects of ascertainment bias on the inference of genetic structure by conducting PCAs using the R package pcadapt (Privé *et al.*, 2020). We performed PCAs using SNPs separately from each source and retained the results from two sets where we observed the strongest effects of ascertainment bias, from sources UOULU RNA-seq and UOULU exomeFEB2019, and one in which the effects were minimal, the ProCoGen haploid sources. To further illustrate the root cause of the observed biases, we performed those PCAs with and without the individuals present in the original discovery panel and driving the patterns observed.

### Linkage map position of PiSy50k markers

To assess whether markers from the PiSy50k array are homogeneously distributed across all chromosomes, we positioned them on a genetic map produced for *P. taeda* by Westbrook *et al*. (2015) comprised of 12 linkage groups (LG) and to which contigs from *P. taeda* reference genome Pita v1.01 have been mapped. We included all PiSy50k SNPs previously mapped to one of the contigs or scaffolds from the same reference genome (data sources ProCoGen haploid and diploid). When a given SNP was outside the aligned segment of the reference contig, we used the closest position effectively aligned on the genetic map from the same contig as a reference point to infer the position of the focal SNP on the map, assuming that the physical distances covered by single contigs from the Pita v1.01 reference genome to be negligible compared to the size of each individual LG.

### PiSy50k array genotyping

We tested the PiSy50k array performance by genotyping 2 688 samples (across seven plates). The 2688 samples consisted of 317 Finnish plus trees, 1847 full-sib offspring from the Finnish breeding population, 489 Scottish samples, three Austrian samples, 11 Estonian samples and 21 controls. The needle control was a single tree from Scotland, UK, and was included on each genotyping plate; this sample had also been genotyped on the Axiom_PineGAP array. In addition, seven haploid megagametophyte samples were genotyped twice, such that each sample was genotyped on two different random plates. Other samples were randomized over the plates such that the different geographic locations and sample categories (plus trees and offspring) were spread on all plates to avoid plate effects that may bias genotyping results of a specific sample category.

The arrays were manufactured by Thermo Fisher Scientific (Waltham, MA, US) and genotyping was conducted by University of Bristol Genomics Facility (Bristol, UK). Needle samples (n = 2 674, including 7 controls) were dried and stored in bags with silica gel. For megagametophyte samples (7 control samples included twice each), germination was initiated by placing the seeds on a moist filter paper inside a petri dish for 24 hours at room temperature. Seeds were then dissected under a microscope to separate megagametophyte from the embryo tissue. The DNA from Finnish and Estonian samples was extracted using E.Z.N.A.® SP Plant DNA Kit (Omega Bio-tek, Inc.). DNA of Scottish needles were extracted using a Qiagen DNeasy Plant kit and checked visually on a 1 % agarose gel. DNA was quantified with a Qubit spectrophotometer.

We performed the genotype call using Axiom Analysis Suite (version 5.1.1.1) following the Axiom Best Practices Workflow with default parameters concordantly to the screening array genotype calling, except for the plate QC threshold for average call rate for passing samples, which we set to 0.97. We retained the markers in the PHR and NMH conversion classes for analyses.

### Evaluation of the PiSy50k array performance

#### Error rate and heterozygosity in haploid samples

We genotyped 21 control samples to estimate error rates for each array: one needle and two megagametophyte controls per plate, with replicate megagametophyte pairs arranged over sequential plates. We estimated the error rates as the proportion of calls which did not match among pairs of controls across plates (excluding calls where one or both were missing). We also measured the heterozygosity in megagametophyte samples to assess probe specificity and identify putative paralogous markers in the PiSy50k array.

#### Pedigree inference and mendelian error rate

We used a subset of 153 samples from 10 crosses, including 18 parents and their 135 offspring, to estimate the coefficients of kinship (K) and the proportion of sites where individuals share no allele (IBS0) between all pairs using converted SNPs (40 405) with KING v2.2.5 (options -- related --degree 2) (Manichaikul *et al*., 2010). We estimated the Mendelian error rate within each family independently using PLINK v1.90b5.2 (option --mendel).

#### Population clustering and ascertainment bias

To evaluate the power of the PiSy50k in discriminating samples from different origins, we used a subset of 120 plus-tree samples: 30 samples from Scotland, grouped in four geographic areas, and 30 samples from Southern, Central and Northern Finland each. We assessed the genetic structure by performing three PCAs: using all 120 samples, the 90 Finnish samples or the 30 Scottish samples separately. We used the function *prcomp* (core R, with scaling and centering options enabled) after replacing missing data for a given genotype by the locus’ allele frequency. Finally, to assess the effect of ascertainment bias on the MAF generated with PiSy50k, we compared the MAF distribution of the Finnish subset of 90 plus trees to the one obtained using exome capture data of Scots pine trees published in Tyrmi *et al*. 2020. From the published vcf file, we extracted the data of 42 megagametophyte samples from four Finnish populations (Inari, Kälviä, Kolari and Punkaharju). We then replaced genotypes with depth below 5 with missing data and kept only loci with a minimum call rate of 50%. Finally, to have comparable MAF distributions, we downsampled both distributions to a sample size of 30.

## Author Contributions

Design of the study: AKN, AP, CK, KK, MH, OS, StC, STK, TP. Field and laboratory work: AKN, AP, SaC, STK, TAK, TP, RK, StC. Computational analyses: AKN, AP, CK, DIO, JST, KA, STK, TMM, TP, WW. Initial draft of the manuscript: AKN, CK, TP. Final manuscript: All authors.

## Acknowledgements

This project has received funding from the European Union’s Horizon 2020 research and innovation programme under grant agreement No 773383 (to UOULU, Luke, UKCEH), Academy of Finland, grants 287431, 293819 and 319313 (to T.P.), 307582 (to O.S.) and 307581 (to K.K.), 309978 (to S.T.K.), and NoE EVOLTREE and Metsänjalostussäätiö (to S.T.K.), and by two grants in the UK: GAPII (NE/K012177/1), funded by NERC, and PROTREE (BB/L012243/1), funded jointly by BBSRC, DEFRA, ESRC, the Forestry Commission, NERC and the Scottish Government, under the Tree Health and Plant Biosecurity Initiative.

The authors wish to thank Joan Beaton and Glenn Iason (James Hutton Institute) for providing Scottish needle samples for genotyping.

## Supporting information legends

The following material is included in the supporting information:

- Supporting Figures S1 to S4
- Supporting Tables S1 to S5
- Supporting Methods S1: “Additional steps/details in selecting markers from screening array to PiSy50k array”.
- Supporting Data S1 and S2

### Legends

**Figure S1**. Minor allele frequencies for the Intensive Study Site Punkaharju (southeast Finland) population (N=466) and 56 693 SNPs without missing data in the screening array. The red line illustrates the expected neutral MAF (Tajima, 1989). Note that this figure is identical to Figure 3 but is represented with a logarithmic scale on both the x- and y-axes.

**Figure S2**. Principal component analysis on the screening array data illustrating the ascertainment bias on the observed genetic structure. (a, c, e) Analysis including samples used in SNP discovery panels of each SNP source, discovery individuals are highlighted and labelled, except in e) for clarity. (b, d, f) Analysis excluding samples used in SNP discovery. SNP sources: (a, b) UOULU RNA-seq (48 357 SNPs), (c, d) UOULU exomeFEB2019 (6 137 SNPs) and (e, f) ProCoGen haploid (23 204 SNPs).

**Figure S3**. Hardy-Weinberg equilibrium (HW) test results for the screening array data before filtering (a,b) and for the selected set for the PiSy50k (c,d). (a,c) Q-Q plots comparing the *p* values expected based on the null distribution against the observed *p* values from the exact HW tests of 10 000 random SNPs on the screening array before (a) and after (c) selecting markers for the PiSy50k array. The green line indicates the expected under HW. (b,d) Ternary plots showing the genotype frequencies of 10 000 random SNPs on the screening array before (b) and after (d) selecting markers for the PiSy50k array. Blue and red dots are markers respectively following or deviating significantly from the HW expectations (Chi-square test at alpha level 0.001).

**Figure S4**. Mendelian errors (ME) of the PiSy50k identified in 40 405 SNPs genotyped in 135 trios (10 crosses). (a) Distribution of ME across loci, the red line indicates the mean error rate across loci (0.29%). (b) ME across families (bars at 0 and 1 indicate the number of SNPs with no ME and with ME in only one family).

**Table S1**. Conversion type for markers from each data set in the screening array based on individuals with call rate 97% or above. We included the markers with the PHR and NMH conversion types (in bold) in the selection of markers for the PiSy50k array. PHR = Poly High Resolution, NMH = No Minor Homozygote, MHR = Mono High Resolution, CRBT = Call Rate Below Threshold, OTV = Off-Target Variant. Values in parenthesis are the proportion (per cent) of each conversion type in each data set.

**Table S2**. Number and proportions of markers from each source at different steps of the PiSy50k array design.

**Table S3**. Distribution of PiSy50k markers on *P. taeda* linkage groups (Westbrook *et al*., 2015). **Table S4**. Conversion type for markers from each data set in the PiSy50k array based on individuals with call rate 97% or above. We included the markers with the PHR and NMH conversion types (in bold) in further analyses. Count and proportion (%) of each conversion type is given within each data set.

**Table S5**. Evaluation of the PiSy50k array for the control samples with call rate above 97%. Values before the forward slash indicate estimates obtained from the full PiSy50k array (40 405 SNPs). Values after the forward slash indicate estimates obtained from the subset of SNPs and the needle sample also genotyped by the Axiom_PineGAP array (7 592 SNPs). CR: call rate; Het: heterozygosity. Mean pairwise error rate estimated as percentage of calls among control pairs that were different (excluding markers which had missing data in at least one of the pairs). **Methods S1**. Additional steps/details in selecting markers from screening array to PiSy50k array.

**Data S1**. The metadata for markers included on the PiSy50k array.

**Data S2**. Shared errors across controls identified in the error evaluation of the PiSy50k, see the main text.

