## Supporting Information for "Taming the massive genome of Scots pine with PiSy50k, a new genotyping array for conifer research"

This document includes:

- Supporting figures S1 to S4
- Supporting tables S1 to S5
- Supporting experimental procedure M1: “Additional steps/details in selecting markers from screening array to PiSy50k array”

Other supporting materials for this manuscript include the following:

- Supporting Data S1: the metadata for markers included on the PiSy50k array.
- Supporting Data S2: shared errors across controls identified during the error evaluation of the PiSy50k.

### 1 Supporting figures

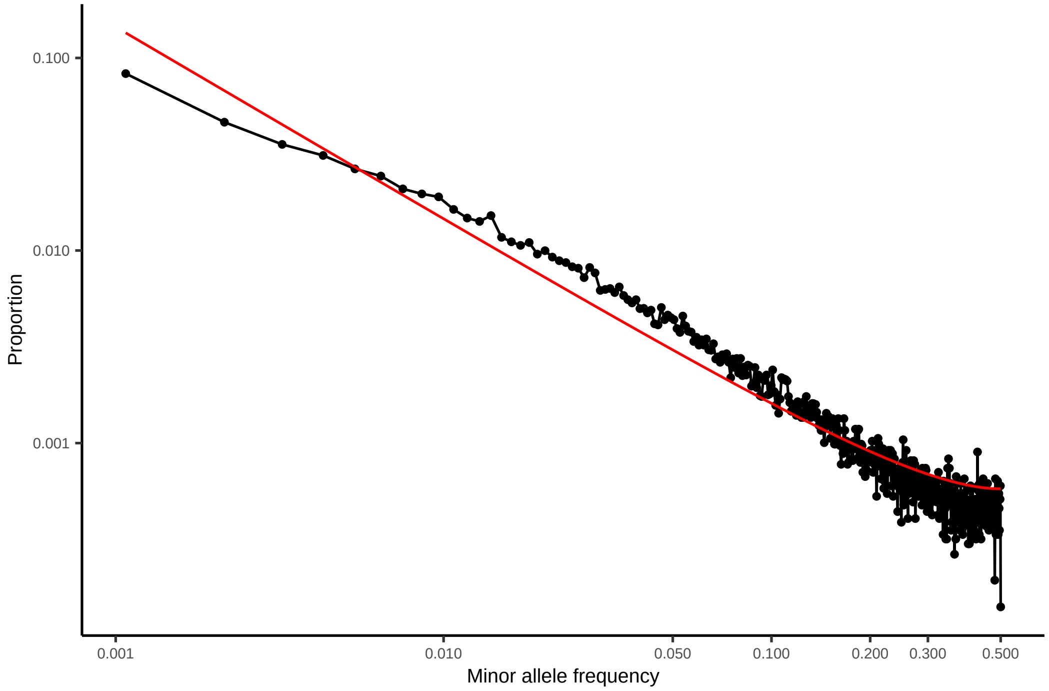


**Figure S1.** Minor allele frequencies for the Intensive Study Site Punkaharju (southeast Finland) population (N=466) and 56 693 SNPs without missing data in the screening array. The red line illustrates the expected neutral MAF (Tajima, 1989). Note that this figure is identical to Figure 3 but is represented with a logarithmic scale on both the x- and y-axes.

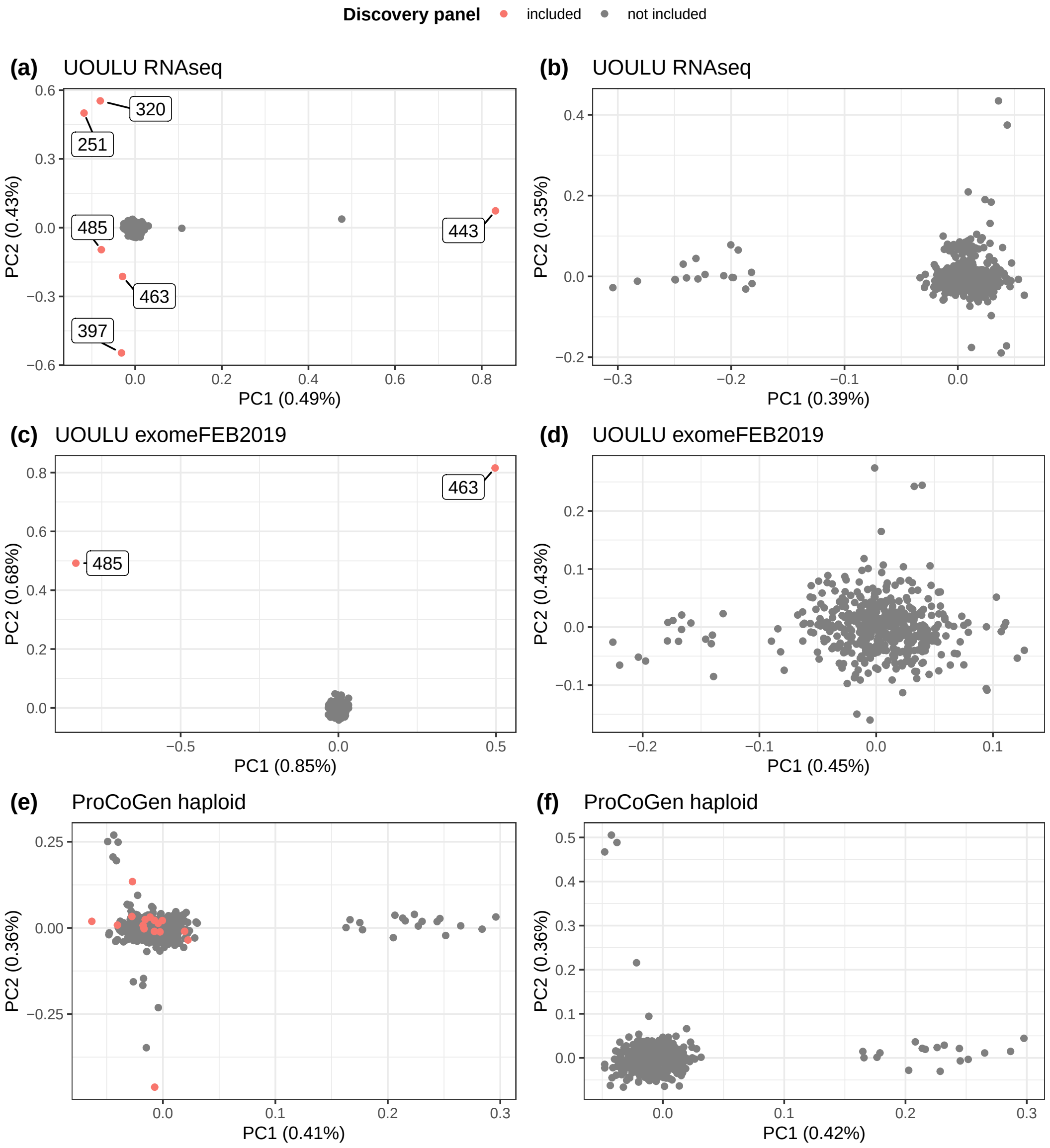


**Figure S2.** Principal component analysis on the screening array data illustrating the ascertainment bias on the observed genetic structure. (a, c, e) Analysis including samples used in SNP discovery panels of each SNP source, discovery individuals are highlighted and labelled, except in e) for clarity. (b, d, f) Analysis excluding samples used in SNP discovery. SNP sources: (a, b) UOULU RNA-seq (48 357 SNPs), (c, d) UOULU exome-FEB2019 (6 137 SNPs) and (e, f) ProCoGen haploid (23 204 SNPs).

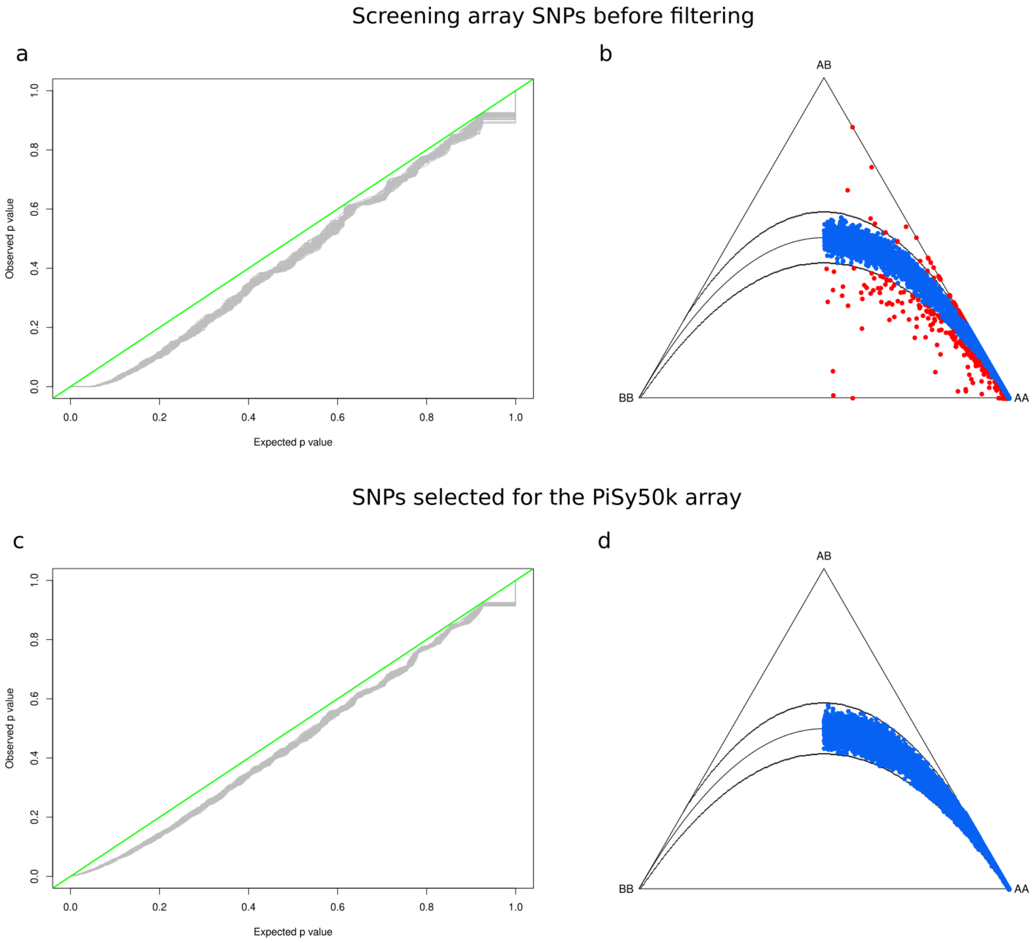


**Figure S3.** Hardy-Weinberg equilibrium (HW) test results for the screening array data before filtering (a,b) and for the selected set for the PiSy50k (c,d). (a,c) Q-Q plots comparing the p values expected based on the null distribution against the observed p values from the exact HW tests of 10 000 random SNPs on the screening array before (a) and after (c) selecting markers for the PiSy50k array. The green line indicates the expected under HW. (b,d) Ternary plots showing the genotype frequencies of 10 000 random SNPs on the screening array before (b) and after (d) selecting markers for the PiSy50k array. Blue and red dots are markers respectively following or deviating significantly from the HW expectations (Chi-square test at alpha level 0.001).

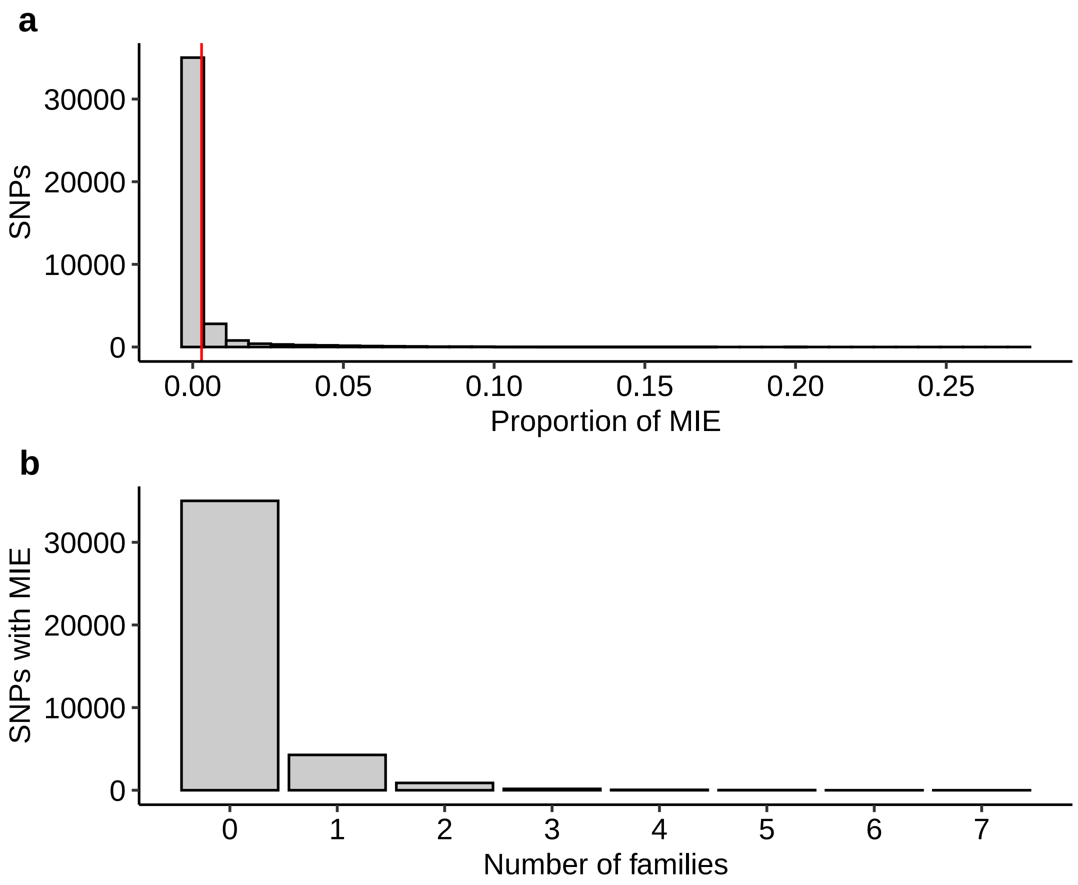


**Figure S4.** Mendelian errors (ME) of the PiSy50k identified in 40 405 SNPs genotyped in 135 trios (10 crosses). (a) Distribution of ME across loci, the red line indicates the mean error rate across loci (0.29%). (b) ME across families (bars at 0 and 1 indicate the number of SNPs with no ME and with ME in only one family).

#### 2 Supporting tables

**Table S1.** Conversion type for markers from each data set in the screening array based on individuals with call rate 97% or above. We included the markers with the PHR and NMH conversion types (in bold) in the selection of markers for the PiSy50k array. PHR = Poly High Resolution, NMH = No Minor Homozygote, MHR = Mono High Resolution, CRBT = Call Rate Below Threshold, OTV = Off-Target Variant. Values in parenthesis are the proportion (per cent) of each conversion type in each data set.

| Data set ID | Nb of markers | PHR | NMH | MHR | CRBT | Other | OTV |
|---|---|---|---|---|---|---|---|
| ProCoGen haploid | 94180 | 16288 (17.3) | 21217 (22.5) | 13527 (14.4) | 5901 (6.3) | 1252 (1.3) | 35995 (38.2) |
| ProCoGen diploid | 23565 | 1395 (5.9) | 7749 (32.9) | 8835 (37.5) | 403 (1.7) | 145 (0.6) | 5038 (21.4) |
| UOULU exomeFEB2019 | 25140 | 6486 (25.8) | 1919 (7.6) | 1871 (7.4) | 2971 (11.8) | 314 (1.2) | 11579 (46.1) |
| UOULU RNA-seq | 179240 | 44592 (24.9) | 23658 (13.2) | 39271 (21.9) | 14801 (8.3) | 4497 (2.5) | 52421 (29.2) |
| UKCEH1 | 18901 | 5750 (30.4) | 3130 (16.6) | 8322 (44) | 280 (1.5) | 207 (1.1) | 1212 (6.4) |
| UKCEH2 | 57261 | 14118 (24.7) | 8765 (15.3) | 12400 (21.7) | 4423 (7.7) | 974 (1.7) | 16581 (29) |
| UOULU candidate | 3273 | 1038 (31.7) | 598 (18.3) | 516 (15.8) | 273 (8.3) | 35 (1.1) | 813 (24.8) |
| LUKE candidate | 5980 | 251 (4.2) | 371 (6.2) | 3082 (51.5) | 256 (4.3) | 117 (2) | 1903 (31.8) |
| Total | **407540** | **89918 (22.1)** | **67407 (16.5)** | 87824 (21.5) | 29308 (7.2) | 7541 (1.9) | 125542 (30.8) |



**Table S2.** Number and proportions of markers from each source at different steps of the PiSy50k array design.

| Data set ID | Initial set | | Screening array | | PiSy50k array | | Screening to PiSy50k array |
|---|---|---|---|---|---|---|---|
| | count | % | count | % | count | % | % |
| ProCoGen haploid | 1870598 | 48.9 | 94180 | 23.1 | 6995 | 14.7 | 7.4 |
| ProCoGen diploid | 304661 | 8 | 23565 | 5.8 | 340 | 0.7 | 1.4 |
| UOULU exomeFEB2019 | 95504 | 2.5 | 25140 | 6.2 | 3356 | 7 | 13.3 |
| UOULU RNA-seq | 1349291 | 35.3 | 179240 | 44 | 20797 | 43.6 | 11.6 |
| UKCEH1 | 20795 | 0.5 | 18901 | 4.6 | 6718 | 14.1 | 35.5 |
| UKCEH2 | 175841 | 4.6 | 57261 | 14.1 | 7788 | 16.3 | 13.6 |
| UOULU candidate | 3584 | 0.1 | 3273 | 0.8 | 1187 | 2.5 | 36.3 |
| LUKE candidate | 6157 | 0.2 | 5980 | 1.5 | 531 | 1.1 | 8.9 |
| Total | 3826431 | 100 | 407540 | 100 | 47712 | 100 | 11.7 |

**Table S3.** Distribution of PiSy50k markers on *P. taeda* linkage groups (Westbrook *et al.* 2015).

| Linkage Group | Length (cM)† | Nb of markers per source (ProCoGen) Haploid | Diploid | Total nb of markers count | % | average distance (cM) |
|---|---|---|---|---|---|---|
| 1 | 184.89 | 141 | 3 | 144 | 8.9 | 3.45 |
| 2 | 222 | 120 | 9 | 129 | 8 | 3.57 |
| 3 | 186.88 | 119 | 4 | 123 | 7.6 | 3.06 |
| 4 | 186.32 | 126 | 8 | 134 | 8.3 | 3.13 |
| 5 | 216.41 | 164 | 5 | 169 | 10.4 | 2.65 |
| 6 | 193.57 | 142 | 6 | 148 | 9.1 | 2.61 |
| 7 | 193.43 | 146 | 11 | 157 | 9.7 | 2.6 |
| 8 | 189.56 | 113 | 5 | 118 | 7.3 | 2.8 |
| 9 | 172 | 126 | 4 | 130 | 8 | 2.34 |
| 10 | 211 | 136 | 9 | 145 | 9 | 2.81 |
| 11 | 146.89 | 128 | 7 | 135 | 8.3 | 2.32 |
| 12 | 202.48 | 83 | 4 | 87 | 5.4 | 3.83 |

†Values taken from Table 2 in Westbrook *et al.* 2015.



**Table S4.** Conversion type for markers from each data set in the PiSy50k array based on individuals with call rate 97% or above. We included the markers with the PHR and NMH conversion types (in bold) in further analyses. Count and proportion (%) of each conversion type is given within each data set.

| Data set ID | Nb of markers on arrays | PHR count | PHR % | NMH count | NMH % | MHR count | MHR % | CRBT count | CRBT % | Other count | Other % | OTV count | OTV % |
|---|---|---|---|---|---|---|---|---|---|---|---|---|---|
| ProCoGen haploid | 6995 | 5244 | 75 | 285 | 4.1 | 3 | 0 | 1043 | 14.9 | 418 | 6 | 2 | 0 |
| ProCoGen diploid | 340 | 255 | 75 | 19 | 5.6 | 2 | 0.6 | 39 | 11.5 | 25 | 7.4 | 0 | 0 |
| UOULU exomeFEB2019 | 3356 | 2682 | 79.9 | 75 | 2.2 | 0 | 0 | 438 | 13.1 | 161 | 4.8 | 0 | 0 |
| UOULU RNA-seq | 20797 | 16788 | 80.7 | 684 | 3.3 | 4 | 0 | 2457 | 11.8 | 860 | 4.1 | 4 | 0 |
| UKCEH1 | 6718 | 4871 | 72.5 | 898 | 13.4 | 652 | 9.7 | 133 | 2 | 131 | 1.9 | 33 | 0.5 |
| UKCEH2 | 7788 | 6541 | 84 | 184 | 2.4 | 0 | 0 | 782 | 10 | 280 | 3.6 | 1 | 0 |
| UOULU candidate | 1187 | 614 | 51.7 | 261 | 22 | 23 | 1.9 | 164 | 13.8 | 123 | 10.4 | 2 | 0.2 |
| LUKE candidate | 531 | 132 | 24.9 | 145 | 27.3 | 43 | 8.1 | 115 | 21.7 | 92 | 17.3 | 4 | 0.8 |
| Total | 47712 | 37127 | 77.8 | 2551 | 5.3 | 727 | 1.5 | 5171 | 10.8 | 2090 | 4.4 | 46 | 0.1 |

**Table S5.** Evaluation of the PiSy50k array for the control samples with call rate above 97%. Values before the forward slash indicate estimates obtained from the full PiSy50k array (40 405 SNPs). Values after the forward slash indicate estimates obtained from the subset of SNPs and the needle sample also genotyped by the Axiom_PineGAP array (7 592 SNPs). CR: call rate; Het: heterozygosity. Mean pairwise error rate estimated as percentage of calls among control pairs that were different (excluding markers which had missing data in at least one of the pairs).

| Tissue | Plate | CR (%) | Het (%) | Mean pairwise error rate (%) |
|---|---|---|---|---|
| Needle | 2 | 98.43 / 98.13 | 29.17 / 24.97 | 1.01 / 0.50 |
| | 3 | 98.52 / 98.34 | 29.30 / 25.11 | 0.98 / 0.56 |
| | 7 | 98.38 / 98.08 | 29.43 / 24.95 | 0.99 / 0.52 |
| Megagametophyte | 4 | 98.66 / — | 00.82 / — | 0.70 / — |
| | 2 | 98.34 / — | 00.97 / — | 0.70 / — |

#### 3 Supporting experimental procedures

##### Additional steps/details in selecting markers from screening array to PiSy50k array

In addition to the steps described in the main text, we performed the following filtering from the screening to the PiSy50k array. During the screening array development, we included multiple probe sets for markers of high piority to be able to select the best performing probe sets. These markers had the same Affy-SNP-ID, but differing probe set sequences. From these, there were 403 markers (806 probe sets) with the conversion type PHR or NMH. We chose the probe sets that were classified as the best probe sets by Thermo Fisher Scientific (BestProbeset = 1). We further selected the best probe sets after filtering for Mendelian errors, heterozygote errors in haploid megagametophyte samples, Hardy Weinberg equilibrium $p$-value, and minor allele frequency, but before filtering for LD. Since we analyzed the screening data before filtering, all 403 duplicate markers were included in the screening array exploration (e.g. Figures 2–3).

Additionally, since we wanted to include as many as possible markers from the previously developed Axiom PineGap array (Perry et al., 2020), we retained all markers with MAF > 0.05 and call rate > 0.8 in a previously genotyped European sample (Perry et al., 2020), as described in the main Material and Methods. During this step, in addition to markers with conversion types PHR and NMH, we also included markers with conversion type MHR on the screening array. All UKCEH1 markers (20 795 markers that performed well on the Axiom PineGAP array) were included in our screening array and shared a probe set id with our screening array markers. All UKCEH1 markers were included in this additional selection step, even if they shared a source with some of our other marker sources (“Data set ID” column of Tables S1 and S2).

We included all high-priority markers from the candidate gene sources (PacBio and UOULU candidate) and from the UKCEH1 source. As a final step to fit a maximal number of markers on the PiSy50k array, we excluded all markers that required allele-specific probes (i.e. SNPs with alleles C/G and A/T) from the low priority sources (ProCoGen haploid, ProCoGen diploid, UKCEH2 and UOULU RNA-seq). Since allele-specific probes take twice the physical space of non-allele-specific probes on the array, including only the latter allowed us to fit more markers on the PiSy50k array in total.
